# A trainable language model for modulating translation rates in non-model organisms by generating upstream untranslated region sequence libraries

**DOI:** 10.64898/2026.04.18.719341

**Authors:** Alexander D. Duggan, Matthew P. Newman, David R. McMillen

## Abstract

Tuning protein expression in non-model organisms is often constrained by the lack of validated genetic parts and predictive design tools. Translational tuning through the modulation of upstream untranslated regions (5′-UTRs) offers a potentially organism-agnostic route, but existing methods typically rely on mechanistic assumptions, prior knowledge that may not be available in non-model contexts, or the screening of sequence libraries. Here, we present a simple generative approach for creating synthetic 5′-UTR libraries based solely on the genomic sequence statistics of any desired organism. The method uses a sliding-window n-gram language model applied to native 5′-UTR sequences to produce novel sequences that preserve organism-specific base distributions and motifs without hard-coding specific motifs or mechanistic rules into inflexible statistical templates.

We have applied this approach to the model bacterium *Escherichia coli* and the non-model probiotic *Limosilactobacillus reuteri*. Libraries of approximately 1,000 sequences were generated for each organism, from which about 100 unique sequences were experimentally tested for translation of a fluorescent reporter protein. In both organisms, the synthetic libraries yielded a broad range of translation levels from this relatively small number of tested variants. Sequences derived from an organism’s own genomic statistics generally performed better in that organism than sequences derived from the other species. Correlations of individual sequence performance across the two species were weak, and thermodynamic predictions of ribosome binding strength showed very little predictive power, especially in the non-model *L. reuteri*.

The results demonstrate that simple statistical language model approaches applied to genomic data can generate functional translational regulatory sequence libraries without detailed mechanistic knowledge or explicit reference to consensus motifs. The approach requires minimal computational resources, avoids reproducing native sequences, and can be readily applied to any organism with a sequenced genome. This strategy may lower technical barriers to expression tuning in non-model organisms.

## Introduction

Tuning the level of protein expression from a cell is a fundamental step in most biological engineering applications: protein expression levels must be tuned to hit desired production rates in bioprocessing; to align with target ligand concentration in biosensing; or to match the required input ranges of other elements in a designed cellular circuit in information processing or control applications. For well-studied organisms such as *Escherichia coli* and *Saccharomyces cerevisiae*, a wide range of methods for tuning protein expression is available, each with known characteristics and an established set of procedures for implementation and troubleshooting; we will call these “model” organisms, following recent usage of that term. Attempts at tuning generally fall into a handful of broad categories [1-4]: (1) transcriptional tuning, in which the promoter of the gene is modified to alter expression; (2) translational tuning, accomplished by modulating the rate at which mRNA transcripts are translated into a protein of interest; and (3) post-translational tuning, in which average functional protein levels are modulated by influencing protein behaviour, for example through the incorporation of peptide tags that alter the degradation or solubility of the protein.

There are compelling reasons to carry out biological engineering in less well-established, non-model organisms: the handful of established usual-suspect organisms represent only a tiny subset of the range of metabolic, genetic, and environmental diversity available in the microbial world, and using the evolved characteristics of an organism can enable access to important properties that are difficult or impossible to reproduce in one of the better established organisms. Non-model organisms may live and function natively in a target environment of interest (such as human or animal intestines, or the soil around plant root systems) and have strong potential to enable the discovery of production pathways for a wide range of compounds through “bioprospecting” and related approaches [5,6].

Working in less well-studied non-model organisms comes at a considerable cost: the standard approaches for tuning protein expression are most often not available. Organisms may not have a known library of transcription factors or transcriptional initiation sequences to be used in transcriptional tuning, they may lack established sets of degradation or solubility peptide tags for post-translational tuning, and protocols and methods may not be sufficiently established to yield consistent results with these methods. This leaves translational tuning as an appealing approach when working in a non-model organism, but this route also often suffers from a lack of validated protocols and/or insufficiently detailed mechanistic information on the regulatory influences involved, which may vary considerably across organisms. Here, we present a simple method of genetic sequence generation and show that it can generate libraries of upstream untranslated regions that exhibit a wide range of translation rates, in both *E. coli* and the non-model organism *Limosilactobacillus reuteri. L. reuteri* is a probiotic intestinal bacterium found in a wide range of vertebrate hosts (including humans), with strain-host specific mucosal adhesion adaptations that favour lengthy gut colonisation; these features are of interest for biological engineering efforts aimed at sensing or manipulating the intestinal environment in humans or other species [7-10].

Immediately before the start codon of a gene of interest is an upstream sequence denoted the 5’-UTR (the UnTranslated Region at the 5’ end of an mRNA transcript), with a functionally relevant length that varies with by organism and genetic context, with eukaryotes generally having longer-range influences than prokaryotes [11,12]; here, we will focus on the 35 bases upstream of the start codon, a suitable length for prokaryotic translational regulation. mRNA transcripts are bound to a ribosome through one or more sequences within the 5’-UTR, with translation into an amino acid chain commencing at the start codon. The details of this ribosomal mRNA capture vary between organisms but generally involve some form of sequence-specific recognition between the mRNA strand and ribosomal RNA (rRNA) sequences incorporated into a ribosomal complex; examples include the Shine-Dalgarno sequence in many bacteria and the family of Kozak sequences in eukaryotes. The process of translational initiation depends in a complex manner on multiple factors, including the secondary structure of the mRNA strand, the nature of the coding sequence itself, and potential influences from the 3’-UTR downstream of the gene of interest [12,13].

The overall translation rate of a given mRNA transcript is a function of these sequence-dependent factors, with the 5’-UTR sequence clearly playing a substantial role in the process [12,13]. Efforts have been made to create translation rate models capable of forward prediction (given a sequence, predict its translation rate) and inverse prediction (for a desired translation rate, generate sequences to yield that rate). Methods have varied, including mechanistic representations of the mRNA-ribosome binding and its associated thermodynamic properties [14,15], kinetic models of the translational initiation process [16,17], and data-driven models that use measured translation levels to extract regularities from the associated sequences and thus enable statistical prediction of the translation rates of novel sequences [18-22]. The approaches most similar to the approach we propose here employ genomic sequence information to generate results in a variety of applications, including the identification of genes or other sequences of interest through hidden Markov models or related statistical approaches [23-25] and the generation of statistics-preserving DNA sequences driven by finite genomic contexts [26]. Though some success has been achieved in these modelling efforts, the ability to reliably obtain desired translation rates on demand remains somewhat elusive. If the requirement is generating a specific translation rate, or covering a desired range of translation rates, in an arbitrary organism that may be a non-model species, we are not aware of a clear predictive solution available that reliably operates across both model to non-model organisms.

Characterizing sets of candidate sequences experimentally offers a strong alternative or complement to predictive sequence generation: libraries of 5’-UTR candidate sequences are synthesized and their translational efficiencies are then measured directly. These approaches have most often been applied in model organisms [13,27,28], but there is no barrier in principle to extending them into non-model organism applications, except perhaps that some screening approaches are based on very large library sizes [21,22], making them challenging to apply without access to substantial infrastructure.

To support our own work in non-model organisms [29,30], we sought to develop a solution with several key properties: (1) it should be as organism-agnostic as possible, allowing the same approach to be applied to any organism (model or non-model); (2) it should not rely on embedded assumptions about the relative influence of portions of the 5’-UTR on the translation process (since such assumptions may fail to transfer between organisms); (3) it should not require intensive *in cellulo* screening of huge libraries of candidate sequences (which may not be practical or affordable in all research contexts); (4) it should not rely on exact replication of existing sequences, to avoid spontaneous homologous recombination, cross-talk, or other forms of interaction between synthetic and genomic sequences; and (5) it should be implementable without recourse to large amounts of computational power.

The solution we present here combines data-driven and library screening approaches, and satisfies all of our goals: (1) it can be applied to any organism with a known genomic sequence; (2) it incorporates no prior assumptions about specific sequence regions; (3) our tests show that generating on the order of 1000 sequences and testing on the order of 100 of them can yield a wide range of translation levels in two different organisms (one model, one non-model); (4) the sequences generated in our approach have a vanishingly small probability of matching any existing sequence from the genome; and (5) the algorithm is straightforward enough to be readily implemented with commercial desktop/laptop levels of computational power.

## Results and Discussion

The method is summarized in Fig 1. In any organism with a sequenced and annotated genome, we initialize our sequence generation with a two-base pair representing the second and third bases of the start codon (most often the TG of an ATG start codon, with rare exceptions). Querying all 5’-UTRs for each annotated gene, we find all instances where this two-base sequence occurs, and determine the fraction of time each of the four possible bases occur in the 0 position (the first base of the start codon). A new base is chosen by generating a random number and selecting the bases with probabilities equal to their fractional representation in the genome. The “n-gram window” of two bases then takes one step upstream, identifying a new pair of bases and initiating a new query into the genome for the fractions of bases upstream of that pair, to be used for the weighted probabilities of each base occurring at the -1 position. Iterating the process of sliding the window upstream generates a sequence of bases with frequencies determined by the organism’s genome, but without any motif-based or mechanistic information explicitly incorporated into the selection; this is equivalent to the way machine learning-based language models step through sequences of words to predict or generate sentences. Resetting the process and generating a new set of pseudo-random numbers will yield a new sequence, and this process can be quickly repeated to generate a library of any desired size.

**Figure 1.**
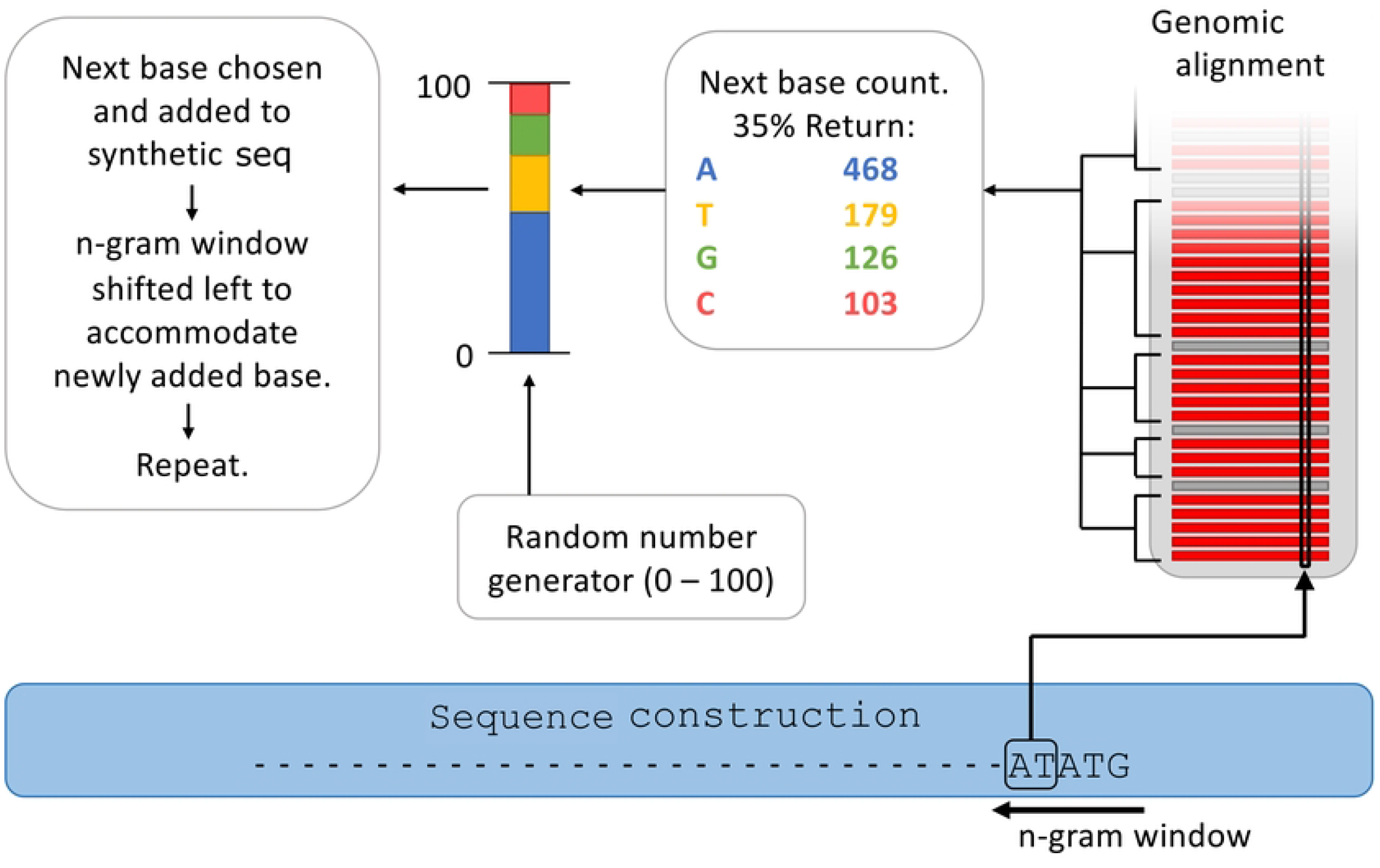
Graphical depiction of the language model. Beginning at the start codon, the algorithm takes the 5’ end of the emerging sequence and queries the genomic 5’ UTR pool for sequences matching the current context. The frequency of each base’s appearance in the set of matching genome sequences sets its probability of being selected in the next position. The selected base is added to the sequence and the n-gram window is shifted one step to the left and used to generate a new context for matching genomic sequences and a new set of per-base probabilities. The cycle repeats until the sequence reaches a desired length; here, all sequences were extended 35 bases upstream of the start codon.

(Generating 10,000 sequences took approximately 12 seconds on a single-core 3.4 GHz CPU.) It is possible to use an n-gram window with n greater than 2, but this comes with a clear trade-off between increasing contextual information (larger n values) and decreasing genomic information.

Larger n-gram windows return smaller sets of matching sequences from the genome of interest (by approximately a factor of 4 for every integer increase in n), and for large enough values of n this can lead to situations where only a single matching sequence is returned from the genome, after which only extensions of that precise sequence will be generated by the algorithm since the frequency of each successive base will be 100% at each step. Our *E. coli* reference genome provides 4459 known 5’-UTR sequences, while our *L. reuteri* reference genome provides 1952 known sequences. A value of n=2 was found to provide an optimal balance between contextual information and sample size, with *L. reuteri* genome queries generally returning numbers of sequences in the low hundreds; increasing to n=3 would reduce this to a few dozen sequences, which we considered to be too small a set to provide robust frequency information for selecting the next base in a candidate sequence without overfitting to a small sample.

We have explored the range of translational efficiencies resulting from varying the 5’ untranslated regions in two bacterial species: *E. coli* (an extremely well-studied microbe); and *L. reuteri* DSM20016 (a non-model microbe that has been the subject of much less extensive study and characterization). Fig 2 shows the frequencies of single bases at each position from +2 to -35 upstream of the start codon, based on examining 4449 5’-UTR sequences from the *E. coli* DH10β genome (NCBI accession number NC_010473), and examining 1952 5’-UTR sequences from the significantly smaller *L. reuteri* DSM20016 genome (strain F275). At the right side of the plots, the dominant ATG start codon appears as near-100% frequencies for A, T, and G in positions 0, 1, and 2; this applies to both organisms, as expected. The enrichment of adenine (A) and guanine (G) bases in the -12 to -7 region in *E. coli* (Fig. 2A) is consistent with the frequent inclusion of a Shine-Delgarno consensus sequence (AGGAGG) in that region. Results from the *L. reuteri* genome show a wider region of stronger G and A enrichment, located slightly further upstream than the equivalent region in *E. coli* (Fig. 2C). The *L. reuteri* frequencies also indicate broad enrichment of adenine across the entire region upstream of the G- and A-enrichment site in line with the organism’s genome-wide low GC content of 38.9% [31], versus the *E. coli* genome’s more even distribution of base frequencies in the equivalent region, in line with its GC content of 50.8% [32].

**Figure 2:**
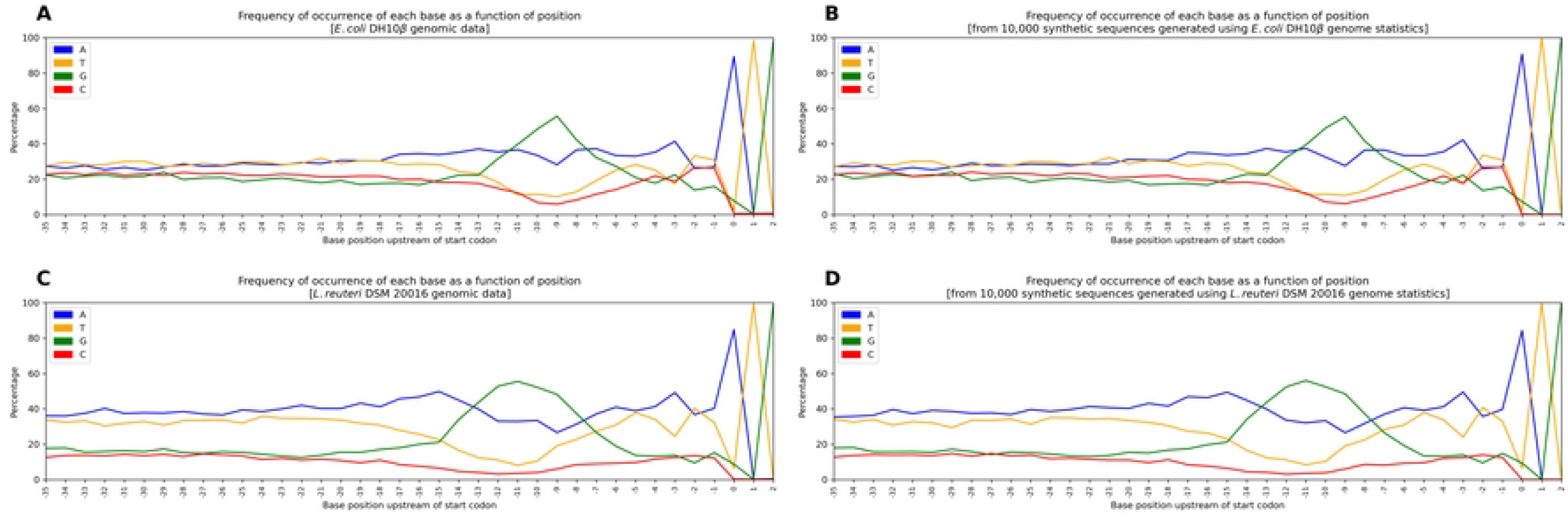
Single base frequencies in the 5’-UTR, in two organismal genomes and in 10,000 sequences generated by our algorithm, from the start codon to position -35 upstream. **(A, B)** The single base frequencies in the *E. coli* DH10β genome (A) and in our algorithmically generated set of sequences (B). **(C, D)** The single base frequencies in the *L. reuteri* DSM20016 genome (C) and in our algorithmically generated set of sequences (D). As expected, the algorithmic sequences retain nearly identical base frequencies, and include features such as the shifted location of a G-rich region between the two organisms, and the overall higher occurrence of adenines in *L. reuteri*’s upstream region versus *E. coli*’s more even distribution among the four bases.

### *in cellulo* testing

We used the approach shown in Fig 1, with n=2, to randomly generate 1000 5’-UTR sequences derived from the genomic statistics of *E. coli* DH10β and *L. reuteri* DSM20016. After plasmid assembly and quality control we were left with over a hundred verified sequences drawn from each organism’s genomic information, and we experimentally tested each sequence in both organisms, finding a wide range of translation levels even in this small library and noting interesting organism-specific properties of the sets of candidate sequences.

5’-UTR sequences generated by the Fig 1 algorithm based on both *E. coli* and *L. reuteri* genomic statistics were incorporated into otherwise identical PTRKH3 plasmid backbones, and transformed into *E* .*coli*. 176 colonies from each RBS pool were picked at random, grown and tested in *E. coli* for activity, and sent for sequencing. After removing duplicate/triplicate 5’-UTR sequences that arose during the assembly process, 123 unique *E. coli* sequences and 117 unique *L. reuteri* sequences remained. These were then transformed into *L. reuteri* DSM 116333 (a naturally derived sub-strain of DSM 20016 with higher transformation efficiency and more stable reporter protein production) [29,30] and tested for activity in the form of fluorescent reporter protein expression. The full sequences lists and experimental measurements are provided as CSV files in Table S1 (*E. coli*-derived sequences) and Table S2 (*L. reuteri*-derived sequences).

Sequences derived from both organisms’ genomic data performed better on average in *E. coli*, yielding both higher average activity and wider activity ranges (Fig 3A, 3B). The *L. reuteri*-derived sequences provided a higher average expression level in *E. coli* than the native *E. coli*-derived sequences did, a feature that is somewhat surprising at first sight. Note, however, that the native upstream sequences in a given organism are not necessarily evolved to *maximize* protein expression levels, but rather to *optimize* them in the full organismal context; it is possible that non-native upstream sequences could shift expression levels higher than an organism’s native average levels.

**Figure 3.**
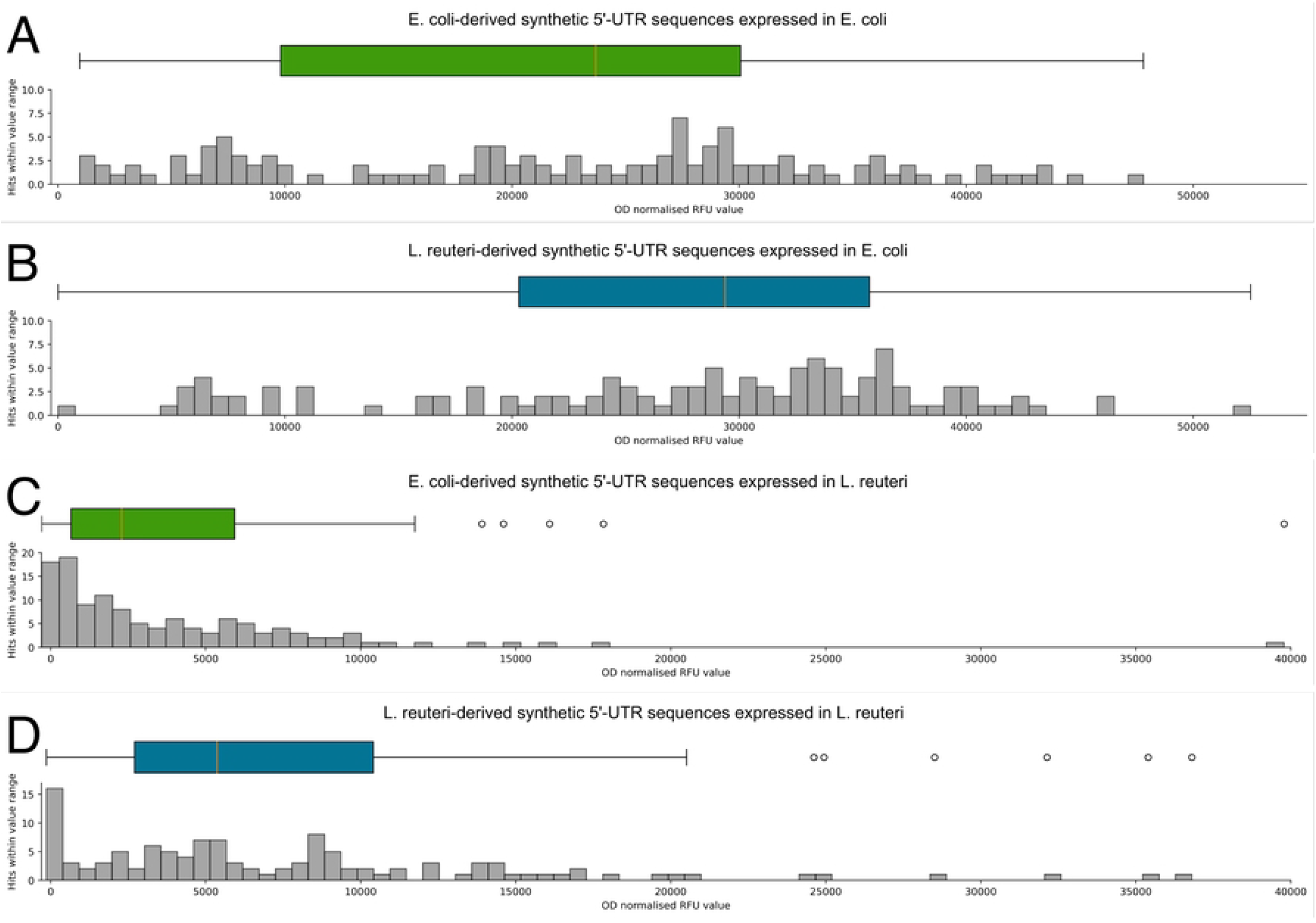
**A)** *E. coli* derived synthetic 5’-UTR sequences transformed into *E. coli* DH10β. **B)** *L. reuteri* derived synthetic 5’ UTR sequences transformed into *E. coli* DH10β **C)** The same *E. coli* derived 5’ UTR sequences from panel A, transformed into *L. reuteri* DSM 116333. **D)** The same *L. reuteri* derived synthetic 5’ UTR sequences from panel B, transformed into *L. reuteri* DSM 116333.

In the non-model organism *L. reuteri*, the *E. coli*-derived synthetic 5’-UTR sequence pool did not yield the features we sought, with significantly lower average activity, many more sequences yielding only minimal activity, and fewer sequences showing moderate to high activity (Fig 3A vs 3C). The 5’-UTR sequences derived from *L. reuteri*’s genomic statistics performed noticeably better in their native *L. reuteri* host, showing a larger set of sequences offering moderate to high activity (Fig 3D). Activity levels in *L. reuteri* did not match the high average levels observed in *E. coli*, but using sequences customized for *L. reuteri* did expand the list of available sequence options substantially (Fig 3C vs 3D).

Every sequence was tested in both bacterial species, enabling us to examine cross-species correlations: to what extent did high/low performance of a given sequence in one organism correlate with similarly high/low performance in the other? Fig 4 plots the activity of each sequence in *L. reuteri* against the activity of the same sequence in *E. coli*, for the *E. coli*-derived sequences and for the *L. reuteri*-derived sequences. In both cases, the conclusion is both visually and mathematically clear: the correlations are weak. The *E. coli*-derived sequences appear to show a very slight positive trend (Fig 4A), but the R^2^ value of 0.03 makes it clear that noise dominates.

**Figure 4.**
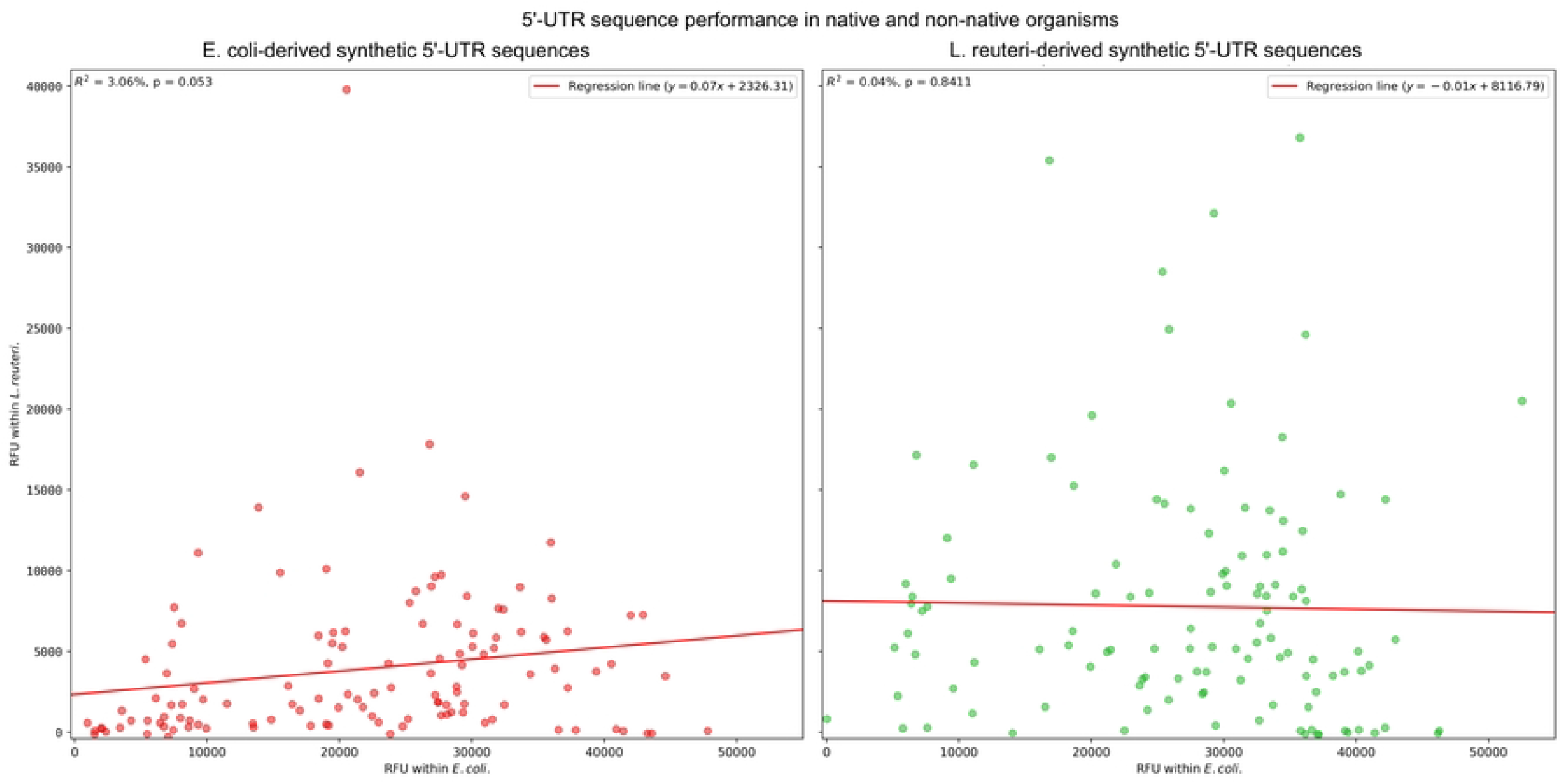
Strengths of our generated 5’-UTR sequences, experimentally measured with fluorescent reporter protein production, in two bacterial species. In each plot, the strength of an individual sequence in *E. coli* is plotted on the horizontal axis, while the strength of the same sequence in *L. reuteri* is plotted on the vertical axis. **(left)** Strengths of 5’-UTR sequences derived from the *E. coli* genome, in the two species; **(right)** Strengths of RBS sequences derived from the *L. reuteri* genome, in the two species. RFU: relative fluorescence units.

*L. reuteri*-derived sequences show a “trend” line that is nearly flat (Fig 4B), and the R^2^ value of 4x10^-4^ makes it clear that there is essentially no relationship between how a given sequence from that pool will perform in the two organisms.

Tools are available to generate predictions of binding strengths between ribosomal 16S subunits and target mRNA sequences and the resulting translational initiation rates, and we have used one well-established tool of this type [14,33-35] to compare these predicted binding strength values for each 5’-UTR sequence in each of our two organism-specific pools to our experimental observations for the corresponding sequence in each organism. Fig 5 shows the result of plotting these predictions against the observed activity in each organism. In *E. coli* (Fig 5A), there is a weak but noticeably positive trend with an R^2^ value of 0.096. In *L. reuteri* (Fig 5B), once again the “trend” line is nearly flat, with the R^2^ value of 5.7x10^-4^ confirming that there is essentially no correlation between the predicted binding strengths and our observed activity levels in the non-model organism.

**Figure 5.**
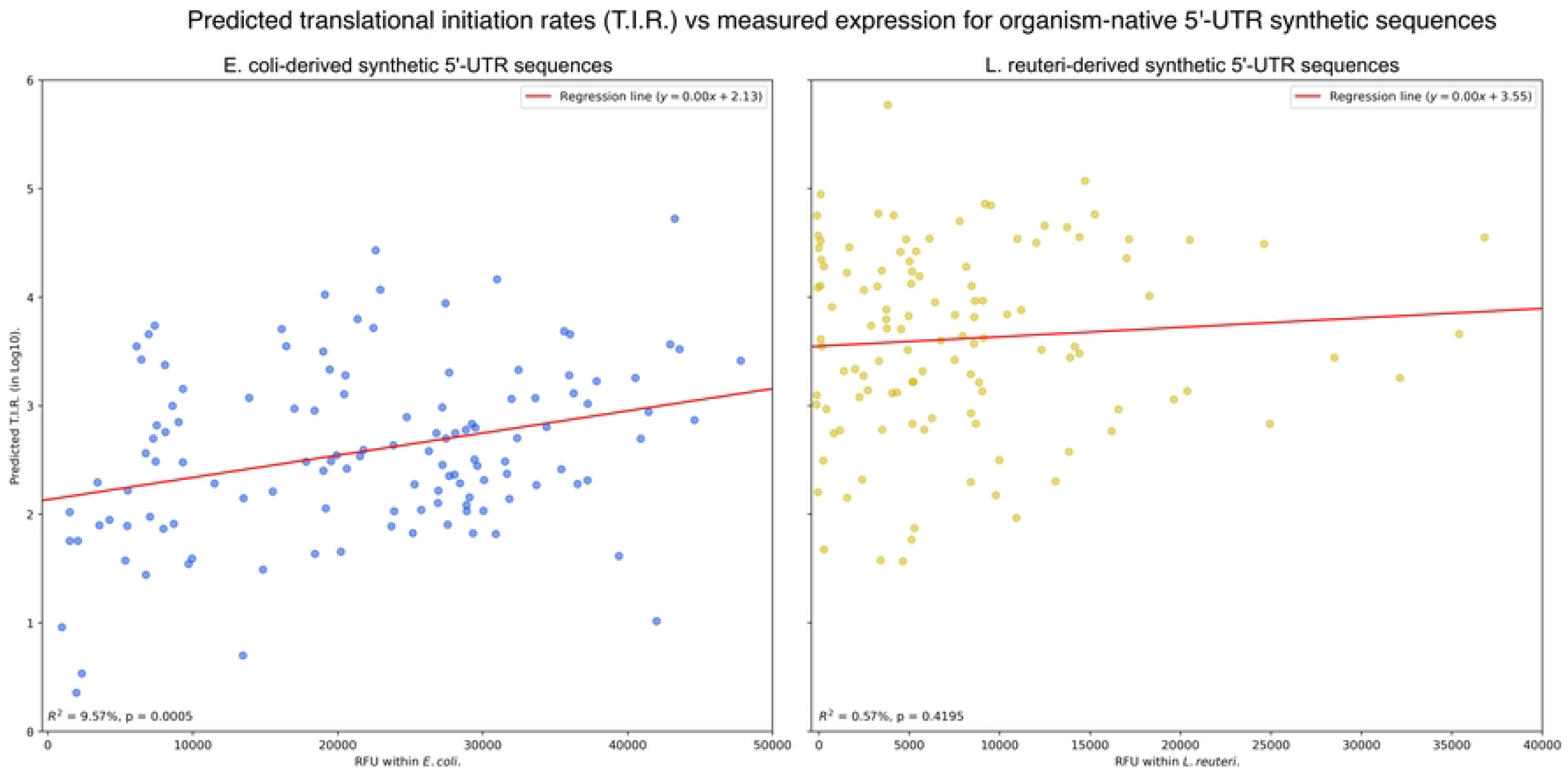
Predicted strengths of our generated 5’-UTR sequences [14,33-35] compared to strengths experimentally measured with fluorescent reporter protein production, in two bacterial species. In each plot, the predicted translational initiation rate (T.I.R.) of sequences derived from the genome of each species is plotted (log_10_ scale) on he vertical axis, against the experimentally measured expression level on the horizontal axis. **(left)** Predicted trengths of 5’-UTR sequences derived from the *E. coli* genome vs measured strengths in *E. coli*. **(right)** Predicted trengths of 5’-UTR sequences derived from the *L. reuteri* genome vs measured strengths in *L. reuteri*. RFU: elative fluorescence units.

Figs 3, 4, and 5 combine to tell a clear story: the model organism *E. coli* is easier to work with across the board than the non-model *L. reuteri*, showing a wider range of translational activity from 5’-UTR sequences derived from its own genomic statistics, as well as having a wider activity range from 5’-UTR sequences derived from the genomic statistics of *L. reuteri*. Individual sequence results from the *E. coli* genome correlated slightly better across the two organisms, and were somewhat more predictable from binding strength estimates. This level of ease of use and predictability is, presumably, correlated with *E. coli*’s status as a highly favoured and widely adopted model organism. *L. reuteri* was generally more difficult to work with: the tested 5’-UTR sequences showed lower overall activity levels (promisingly increased when using statistics from its own genome) with fewer sequence options available to span a wide range of activities, nearly non-existent correlations in sequence performance across the two organisms, and effectively no correlation between predicted binding strengths and experimental activity levels. It is plausible that these features may be related to *L. reuteri*’s status as a non-model organism, with lower ease of use and predictability tending to disfavour its use in routine experimental applications.

The simplicity of our approach neglects long-range influences in the genomic sequence space, and does not bring to bear cutting-edge technologies like machine learning approaches or other sophisticated methods for the analysis of large data sets such as complete organismal genome sequences. We would note, however, that this simplicity has its own positive features: the computational demands are minimal, the success rate is high enough that screening even small libraries is likely to yield sequences offering a range of expression levels, and switching between organisms is a simple matter of priming the algorithm with a different genomic sequence. These features position the approach to be widely accessible, lowering the financial and infrastructure barriers that might otherwise hamper investigations in non-model organisms. Though our approach worked better in the model organism we tested, it is less critical in that context: when working in *E. coli*, there is no shortage of well-validated methods available for tuning gene expression at every level, whereas work in *L. reuteri* or other non-model organisms may suffer from a distinct lack of equivalent options.

The results we have obtained here clearly illustrate that even a simplified language model based on a “sliding window” approach to deriving sequence statistics from genomic information can generate a wide range of translation levels from a tractable library size. Further development of the approach could involve its application to new organisms, and to new classes of sequences, such as promoters or origins of replication.

## Methods

### Sequence generation

The code for the sequence generation algorithm is available at: https://github.com/AIex-Duggan/RBS-Language-Model.

5’-UTR sequences were extracted 35 bases upstream of the start codon (including putative start codons) and aligned to the start codon site. The algorithm begins with a 38 base sequence in which any base is possible at any position. The model for this project was fixed to start with the ‘TG’ codons of the regular and alternative start codons and allowed to deduce next positions from that initial seed, extending the sequence from right to left (equivalently: from downstream to upstream, or from the 3’ to 5’ end of the sequence). The model utilises a simple sliding window n-gram design, with n=2 in our current implementation. At each position, the next base in the sequence is selected at random, with the selection probability for each base determined by its frequency in the set of matching sequences from the target organism’s genome, where a genomic sequence matches if it has the same two bases in the equivalent positions relative to the start codon. The probabilities are given by

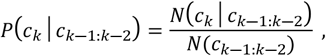

*N*(*c*_*k*−1:*k*−2_) where at position *k*, the probability, *P*, of selecting each base [*c*_*k*_] is set to the fraction of genomic sequences in which it appears (the number, *N*, of such sequences, divided by the total number of matching sequences obtained from the organism’s genome).

### Plasmid construction

Plasmid pTRKH3_mCherry2 (first developed in previous work [29,30] was modified with standard PCR to remove the “universal RBS” BBa_K2918014 sequence, as well as removing the start codon for mCherry2.

Synthetic 5’-UTR sequences generated by the algorithm were augmented at the point of synthesis with additional bases at both ends containing Golden Gate restriction sites to allow scarless integration into a linearized backbone and primer sites to allow PCR amplification. Oligonucleotides were obtained in two batches (one for each organism) containing the full set of DNA sequences in a single-tube mixed pool (Twist Biosciences). Each oligonucleotide pool was subjected to PCR to convert from ssDNA to dsDNA, following a PCR protocol modified from manufacturer recommendations (Twist Biosciences), replacing the recommended KAPA polymerase with Q5 polymerase for improved sequence fidelity. Golden Gate assembly was used to assemble the synthetic 5’-UTR sequences and linear PCR products obtained from the plasmid backbone, and ligation of the final product into the original pTRKH3 backbone placed the 5’-UTR sequences upstream of the mCherry2 gene. See Figure S1 in the Supplementary Information for primer schematics and complete sequences.

#### Cell handling and measurement

The two plasmid pools (with synthetic 5’-UTR sequences derived from *E. coli* and from *L. reuteri*) were each transformed into chemically competent *E. coli* DH10β and plated on LB-agar for isolation. 176 colonies were isolated from each condition, grown in 6 ml LB broth with appropriate selection antibiotic, subject to plasmid extraction (QIAprep mini-prep, Qiagen Canada). All plasmids were sent for sequencing with primers designed to capture the full region of the synthetic sequences and the start of the mCherry2 gene.

The isolates were subjected to several rounds of quality control, eliminating incorrect ligation reaction products, low quality sequencing reads, duplications and suspected contamination. 123 *E. coli* RBS sequences and 117 *L. reuteri* sequences remained in the final pools, which were then transformed into both chemically competent *E. coli* DH10β and electrocompetent *L. reuteri* DSM116333 [29].

*E. coli* DH10β was grown in LB broth (BioShop, Canada) aerobically at 37°C with shaking at 200RPM, LB agar was grown aerobically at 37°C for 16 – 24 hours. Erythromycin was used to maintain pTRKH3 plasmid selection at 250µg/ml in both broth and agar.

*L. reuteri* DSM116333 was grown aerobically in autoclaved MRS broth (Oxoid, UK) at 37°C with no shaking, MRS agar was grown anaerobically at 37°C for 24 – 48 hours or until colonies were visible. Erythromycin was used to maintain pTRKH3 plasmid selection at 10µg/ml in both broth and agar.

For fluorometric assays *E. coli* was grown in LB broth (conditions as above) with selective antibiotic, and after 24 hours 200 µl aliquots were transferred to a flat-bottom optical 96-well plates for OD600 and mCherry2 production measurement (Ex:589 Em:610). *L. reuteri* was grown for 24 hours in filter sterilised MRS broth (conditions as above), and 200 µl aliquots were measured with the same Ex:Em parameters.

## Acknowledgements

A.D. acknowledges David McMillen, Zhe Tang, Matthew Newman, Danny Huong, and Stan Wong for keeping him sane inside the lab, and Ria Lynch for keeping him sane outside it. D.M. is just relieved that A.D. was able to remain sane *anywhere*; if you’ve worked with *L. reuteri*, you may understand.

## Supporting Information captions

**Figure S1**. Synthetic 5′-UTR sequence primers for Golden Gate assembly.

**Table S1**. CSV file containing the complete sequences of the 123 synthetic 5′-UTR sequences derived from *E. coli* genome statistics. Fluorescence measurements for each sequence when tested in both *E. coli* and *L. reuteri*.

**Table S2**. CSV file containing the complete sequences of the 117 synthetic 5′-UTR sequences derived from *L. reuteri* genome statistics. Fluorescence measurements for each sequence when tested in both *E. coli* and *L. reuteri*.

## References

1. Chang, M., Ahn, S.J., Han, T., Yan, D. Gene expression modulation tools for bacterial synthetic biology. Biotechnol Sustain Mater 2024; 1: 6. 10.1186/s44316-024-00005-y.

2. Li, C., Jiang, T., Li, M., Zou, Y., Yan, Y. Fine-tuning gene expression for improved biosynthesis of natural products: From transcriptional to post-translational regulation. Biotechnol. Adv. 2022; 54: 107853. 10.1016/j.biotechadv.2021.107853.

3. Arpino, J.A.J., Hancock, E.J., Anderson, J., Barahona, M., Stan, G.V., Papachristodoulou, A. et al./person-group>. Tuning the dials of synthetic biology. Microbiology 2013; 159: 1236–1253. doi: 10.1099/mic.0.067975-0. Epub 2013 May 23. PMID: 23704788; PMCID: PMC3749727.

4. Ang, J., Harris, E., Hussey B.J., Kil, R., McMillen, D.R. Tuning response curves for synthetic biology. Trends Biotechnol. 2013; 31: 138–146. 10.1021/sb4000564.

5. Heck, M. and Neely, B.A. Proteomics in non-model organisms: A new analytical frontier. J. Proteome Res. 2020; 19: 3595–3606. 10.1021/acs.jproteome.0c00448.

6. Chen, J., Jia, Y., Sun, Y., Kun, L., Zhao, C., Liu, C. et al./person-group>. Global marine microbial diversity and its potential in bioprospecting. Nature 2024; 633: 371–379. 10.1038/s41586-024-07891-2.

7. Moretti, C. H., Grasset, E., Zhe, J., Yang, G., Olofsson, L.E., Tanweer Khan, M. et al. Identification of human gut bacteria that produce bioactive serotonin and promote colonic innervation. Cell Reports 2025; 44: 116434. 10.1016/j.celrep.2025.116434.

8. Alexander, L.M., Khalid, S., Gallego-Lopez, G.M., Astmann, T.J., Oh, J., Heggen, M. et al./person-group>. Development of a Limosilactobacillus reuteri therapeutic delivery platform with reduced colonization potential. Appl Environ Microbiol 2024; 90:e00312–24. 10.1128/aem.00312-24.

9. Poppeliers, J., Boon, M., De Mey, M., Masschelein, J., Lavigne, R. Non-model bacteria as platforms for endogenous gene expression in synthetic biology. Nat Rev Bioeng 2026; 4: 67–81. 10.1038/s44222-025-00354-x.

10. Chan, D.T.C., Bjerg, J., Bernstein, H.C. Broad-host-range synthetic biology: Rethinking microbial chassis as a design variable. ACS Synth Biol. 2025; 14: 3815-3821. https://pubs.acs.org/doi/10.1021/acssynbio.5c00308.

11. Lin, Z., Li, W.H. Evolution of 5’ untranslated region length and gene expression reprogramming in yeasts. Mol Biol Evol. 2012; 29:81–9. 10.1093/molbev/msr143.

12. Zheng, D., Persyn, L., Wang, J., Liu, Y., Ulloa-Montoya, F., Cenik, C. et al./person-group>. Predicting the translation efficiency of messenger RNA in mammalian cells. Nat. Biotechnol. 2025; 10.1038/s41587-025-02712-x.

13. Mutalik, V.K., Guimaraes, J.C., Cambray, G., Mai, Q-A., Christoffersen, M.J., Lance, L. et al./person-group>. Quantitative estimation of activity and quality for collections of functional genetic elements. Nat. Methods 2013; 10: 347–353. 10.1038/nmeth.2403.

14. Salis, H.M., Mirsky, E.A., Voigt, C.A. Automated design of synthetic ribosome binding sites to control protein expression. Nat. Biotechnol. 2009; 27: 946–950. 10.1038/nbt.1568.

15. Roots, C.T., Lukasiewicz, A., Barrick J.E. OSTIR: open source translation initiation rate prediction. J. Open Source Softw. 2021; 6: 3362. 10.21105/joss.03362.

16. Na, D., Lee, D. RBSDesigner: software for designing synthetic ribosome binding sites that yields a desired level of protein expression, Bioinformatics 2010; 26: 2633–2634. 10.1093/bioinformatics/btq458.

17. Na, D., Lee, S., Lee, D. Mathematical modeling of translation initiation for the estimation of its efficiency to computationally design mRNA sequences with desired expression levels in prokaryotes. BMC Syst. Biol. 2010; 4: 71. 10.1186/1752-0509-4-71.

18. Tietze, L., Lale, R. Importance of the 5′ regulatory region to bacterial synthetic biology applications. Microb. Biotechnol. 2021; 14: 1751–7915. 10.1111/1751-7915.13868.

19. Gilliot, P.A., Gorochowski, T.E. Transfer learning for cross-context prediction of protein expression from 5’UTR sequence. Nucleic Acids Res. 2024; 52:e58. 10.1093/nar/gkae491.

20. Terai, G., Asai, K. Improving the prediction accuracy of protein abundance in Escherichia coli using mRNA accessibility. Nucleic Acids Res. 2020; 48: e81. 10.1093/nar/gkaa481.

21. Cuperus, J.T., Groves, B., Kuchina, A., Rosenberg, A.B., Jojic, N., Fields, S. et al./person-group>. Deep learning of the regulatory grammar of yeast 5’ untranslated regions from 500,000 random sequences. Genome Res. 2017; 27: 2015–2024. 10.1101/gr.224964.117.

22. Cambray, G., Guimaraes, J.C., Arkin, A.P. Evaluation of 244,000 synthetic sequences reveals design principles to optimize translation in Escherichia coli. Nat. Biotechnol. 2018; 36: 1005–1015. 10.1038/nbt.4238.

23. Ma, Y., Chen, H., Kang, J., Guo, X., Sun, C., Xu, J., et al. The hidden Markov model and its applications in bioinformatics analysis. Genes & Diseases 2026; 13: 101729. 10.1016/j.gendis.2025.101729.

24. Krogh, A., Mian, I.S., Haussler, D. A hidden Markov model that finds genes in E. coli DNA. Nucleic Acids Res. 1994. 22: 4768–4778. 10.1093/nar/22.22.4768.

25. Salzberg, S.L., Delcher, A.L., Kasif, S., White, O. Microbial gene identification using interpolated Markov models. Nucleic Acids Res. 1998; 26: 544–548. 10.1093/nar/26.2.544.

26. Pinho, A.J., Neves, A.J.R., Martins, D.A., Bastos, C.A.C., Ferreira, P.J.S.G. Finite-context models for DNA coding. In: Miron, S., editor. Signal Processing. InTech [online], pp 117–130. 10.5772/3472.

27. Mutalik, V.K., Guimaraes, J.C., Cambray, G., Lam, C., Christoffersen, M.J., Mai, Q-A. et al./person-group>. Precise and reliable gene expression via standard transcription and translation initiation elements. Nat. Meth. 2013; 10: 354–360. 10.1038/nmeth.2404.

28. Kosuri, S., Goodman, D.B., Cambray, G., Mutalik, V.K., Gao, Y. Arkin, A.P. et al. Composability of regulatory sequences controlling transcription and translation in Escherichia coli, Proc. Natl. Acad. Sci. U.S.A. 2013; 110: 14024–14029. 10.1073/pnas.1301301110.

29. Duggan, A.D., Dillon, M.M., McMillen, D.R.. A promising novel strain of L. reuteri DSM20016 as a chassis for synthetic biology applications. Frontiers Synth. Biol. 2025; 3: 1473338. 10.3389/fsybi.2025.1473338.

30. Duggan, A.D., McMillen, D.R.. Methods for electroporation and transformation confirmation in Limosilactobacillus reuteri DSM20016. J. Visualized Experiments 2023; 196: e65463. 10.3791/65463.

31. Kim, D., Cho, M.J., Cho, S., Lee, Y., Byun, S.J., Lee, S. Complete genome sequence of Lactobacillus reuteri Byun-re-01, isolated from mouse small intestine. Microbiol. Resour. Announc. 2018; 7: e00984–18. 10.1128/MRA.00984-18.

32. Durfee, T., Nelson, R., Baldwin, S., Plunkett, G., Burland, V., Mau, B. et al./person-group>. The complete genome sequence of Escherichia coli DH10B: Insights into the biology of a laboratory workhorse. J. Bacteriol. 2008; 190: 2597–2606. 10.1128/jb.01695-07.

33. Cetnar, D.P., Salis, H.M. Systematic quantification of sequence and structural determinants controlling mRNA stability in bacterial operons. ACS Synth. Biol. 2021; 10: 318–332. 10.1021/acssynbio.0c00471.

34. Reis, A.C., Salis, H.M. An automated model test system for systematic development and improvement of gene expression models. ACS Synth. Biol. 2020; 9: 3145–3156. 10.1021/acssynbio.0c00394.

35. Espah Borujeni, A., Cetnar, D., Farasat, I., Smith, A., Lundgren, N., Salis, H.M. Precise quantification of translation inhibition by mRNA structures that overlap with the ribosomal footprint in N-terminal coding sequences. Nucleic Acids Res. 2017; 45: 5437–5448. 10.1093/nar/gkx061.

